# Organoid single-cell profiling identifies a transcriptional signature of glomerular disease

**DOI:** 10.1101/468850

**Authors:** Jennifer L. Harder, Rajasree Menon, Edgar A. Otto, Jian Zhou, Sean Eddy, Noel L. Wys, Viji Nair, Cristina Cebrian, Jason R. Spence, Olga G. Troyanskaya, Jeffrey Hodgin, Roger C. Wiggins, Benjamin S. Freedman, Matthias Kretzler, European Renal cDNA Bank, Nephrotic Syndrome Study Network

## Abstract

Podocyte injury is central to many forms of kidney disease, but transcriptional signatures reflecting podocyte injury and compensation mechanisms are challenging to analyze *in vivo*. Human kidney organoids derived from pluripotent stem cells (PSCs), a new model for disease and regeneration, present an opportunity to explore the transcriptional plasticity of podocytes. Here, transcriptional profiling of over 12,000 single cells from human PSC-derived kidney organoid cultures was used to identify robust and reproducible cell-lineage gene expression signatures shared with developing human kidneys based on trajectory analysis. Surprisingly, the gene expression signature characteristic of developing glomerular epithelial cells was also observed in glomerular tissue from a kidney disease cohort. This signature correlated with proteinuria and inverse eGFR, and was confirmed in an independent podocytopathy cohort. Three genes in particular were further identified as critical components of the glomerular disease signature. We conclude that cells in human PSC-derived kidney organoids reliably recapitulate the developmental transcriptional program of podocytes and other cell lineages in the human kidney, and that the early transcriptional profile seen in developing podocytes is reactivated in glomerular disease. Our findings demonstrate an innovative approach to identifying novel molecular programs involved in the pathogenesis of glomerulopathies.

## INTRODUCTION

Elucidating the molecular events that underlay the evolution of human glomerular disease has been challenging. While animal and 2D cell culture models have added significantly to the understanding of podocytopathies (1), they are limited in their ability to accurately reflect events in human kidney disease (2, 3). A particularly pertinent gene expression network within a specific cell type can be masked in bulk tissue transcriptional profiling of diseased human kidney tissue by diluting the signal of interest below level of significance, especially when cells are of low abundance like podocytes. Methods to improve detection of glomerulus-specific gene expression alterations have provided some insight, including transcriptional profiling of microdissected glomeruli from individuals with kidney disease (4-7) and enhancing cell-specific signals from kidney tissue transcriptional analysis by *in silico* nanodissection (deconvolution of expression “signatures” based on known cell-specific markers such as *NPHS2*/podocin) (8). However, decrease in podocyte-specific gene expression either by loss of podocytes or by transcriptional modulation introduces further complexity to the gene expression analysis.

Recent technological advances including PSC-derived kidney organoid cultures (9-12) and single cell RNA sequencing (scRNA-seq) (13, 14) present an opportunity to more precisely explore the podocyte developmental transcriptional program. Human kidney organoids can be derived from PSCs over the course of ~ 15 days of differentiation and represent a simplified cohort of kidney cell types. Cells self-organize *in vitro* in patterns that resemble nephron subunits, and include podocytes that can be clearly discerned from neighboring tubular cells. By enabling transcriptional profiling of individual cell types within this heterogeneous cell population, scRNA-seq of human PSC-derived kidney organoid cultures may improve the ability to identify and investigate molecular events more specific to the human kidney. Use of such organoids could be expanded through more careful definition of both cell types generated and their relevancy to human kidney (10, 15-17), assessment of reproducibility, and limitation of non-kidney cell types in kidney organoid cultures.

In the present study we sought to address these concerns by defining the podocyte transcriptional program generated in human PSC-derived kidney organoids in the context of developing human kidney. By performing a novel type of trajectory analysis using combined single cell transcriptomic datasets, we were able to define the repertoire of cells in organoid cultures and to benchmark them against cell types in the developing human kidney. This led to the discovery of a novel gene signature of an early podocyte lineage in organoids, which was used to interrogate gene expression in tissues from diseased human glomeruli. Our results delineate where human PSC-derived kidney organoid cultures can help to define human kidney disease, provide more specific context to the relative transcriptional “age” of cells produced in kidney organoid cultures relative to the developing human kidney and establish the robustness and reproducibility of this system. In addition to identifying novel molecular programs involved in the pathogenesis of glomerulopathies, these findings provide a foundation for researchers to explore therapeutics aimed at reversing these alterations.

## RESULTS

### Single cell transcriptional profiling reveals two putative podocyte cell clusters in kidney organoids

To gain insight into the transcriptional program of podocytes, we transcriptionally profiled podocytes created in kidney organoids. Organoid cultures were generated using UM77-2 human embryonic stem cells (hESCs) and the matrix sandwiching protocol (9, 16). Kidney organoids appeared as discrete clusters of translucent tubular structures, which could be identified by light microscopy (16). We confirmed that organoids contained cells expressing proteins typical of podocytes as well as proximal and distal tubular epithelial cells in nephron-like segments (Fig. 1A-C).

**Figure 1.**
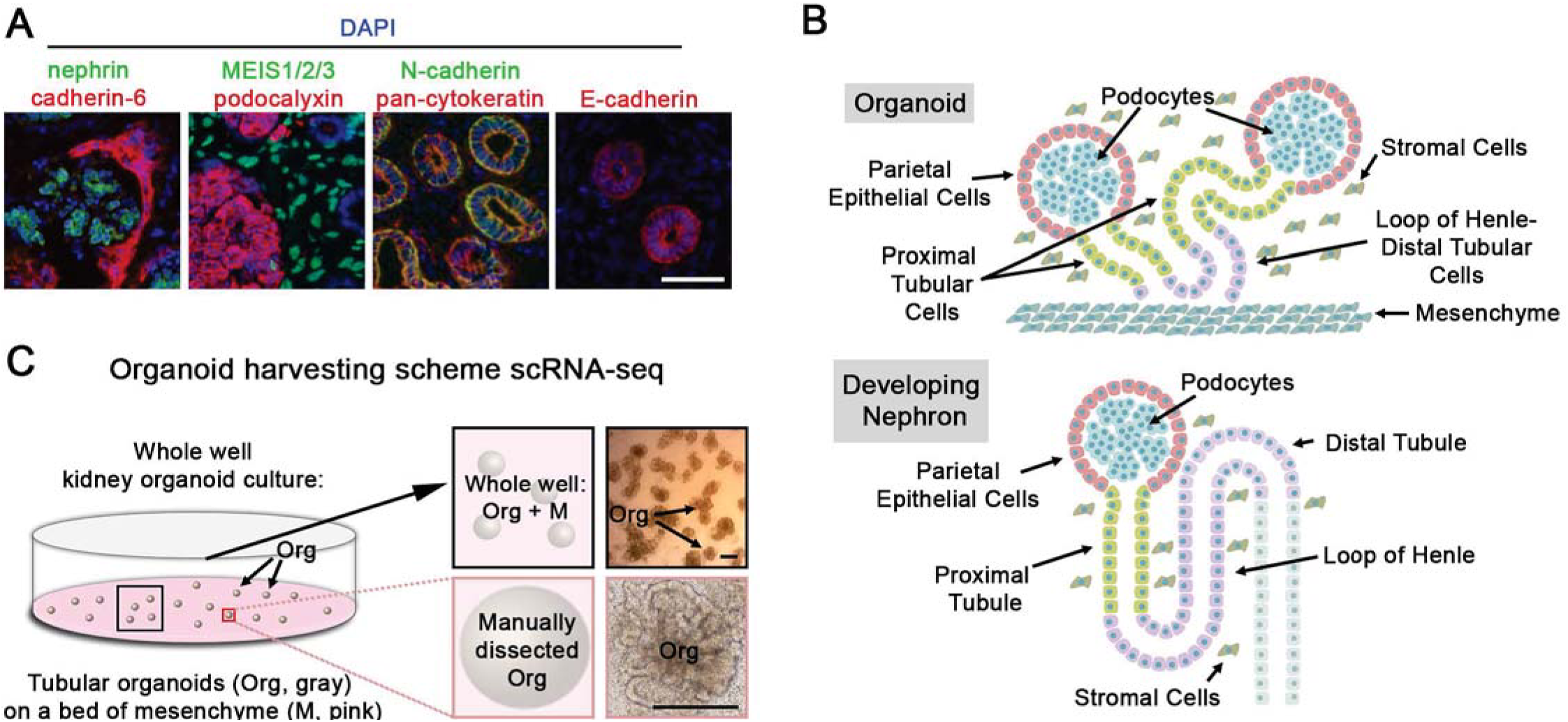
Kidney organoids profiled by single cell transcriptomics contain nascent nephrons. **(A)** Immunofluorescence imaging of sectioned kidney organoids containing nascent podocytes (nephrin, podocalyxin), stromal (MEIS1/2), and epithelialized tubular (Cadherin 6, cytokeratin, N-cadherin, E-cadherin) cells, bar = 100 μm. (**B**) Cartoon showing organoid cultures *in situ* and corresponding developing nephron structures. **(C)** Cartoon depicting organoid harvesting scheme for individual cell analysis by Drop-seq, with corresponding brightfield microscopic images of organoid cultures, black bar = 0.5 mm. Either whole well (organoids plus mesenchyme bed) or manually dissected organoids (20-50) were collected and dissociated per experiment. See related Fig. 2.

To comprehensively characterize cell types present in the cultures and to establish reproducibility of single cell transcriptional profiling of organoids, all cells in the organoid culture wells (“whole well”) were collected (Fig 1C). Analysis of 10,113 single cell transcriptomes from 7 datasets using an unsupervised cell-clustering algorithm generated 11 separate cell clusters (Fig. 2A). Individual clusters contained between 159-1,751 cells and were defined by between 73-272 differentially expressed genes (Fig. S1A). Cells included in this analysis were of consistent high quality with low mitochondrial content (mean 5.1%) and consistent gene number and transcripts, while cell clusters showed good separation by differential gene expression (Figs. S1B-C). The full gene list used to define each cluster is shown in Table S1, while expression profiles of characteristic genes used to identify cell type included in clusters are shown in Fig. 2B. Several of these clusters contained cells that did not express obvious kidney-associated genes and gene expression was more consistent with non-kidney stromal, neural, and proliferating cells (18). However, 6 of the 11 clusters (68.6% of transcriptomes analyzed) could be provisionally assigned as kidney-associated lineage by inspection of each cluster’s gene list (outlined by a dotted line in Fig. 2A), and were selected for viewing in subsequent images (further referred to as “kidney” clusters) (Fig. 2C).

**Figure 2.**
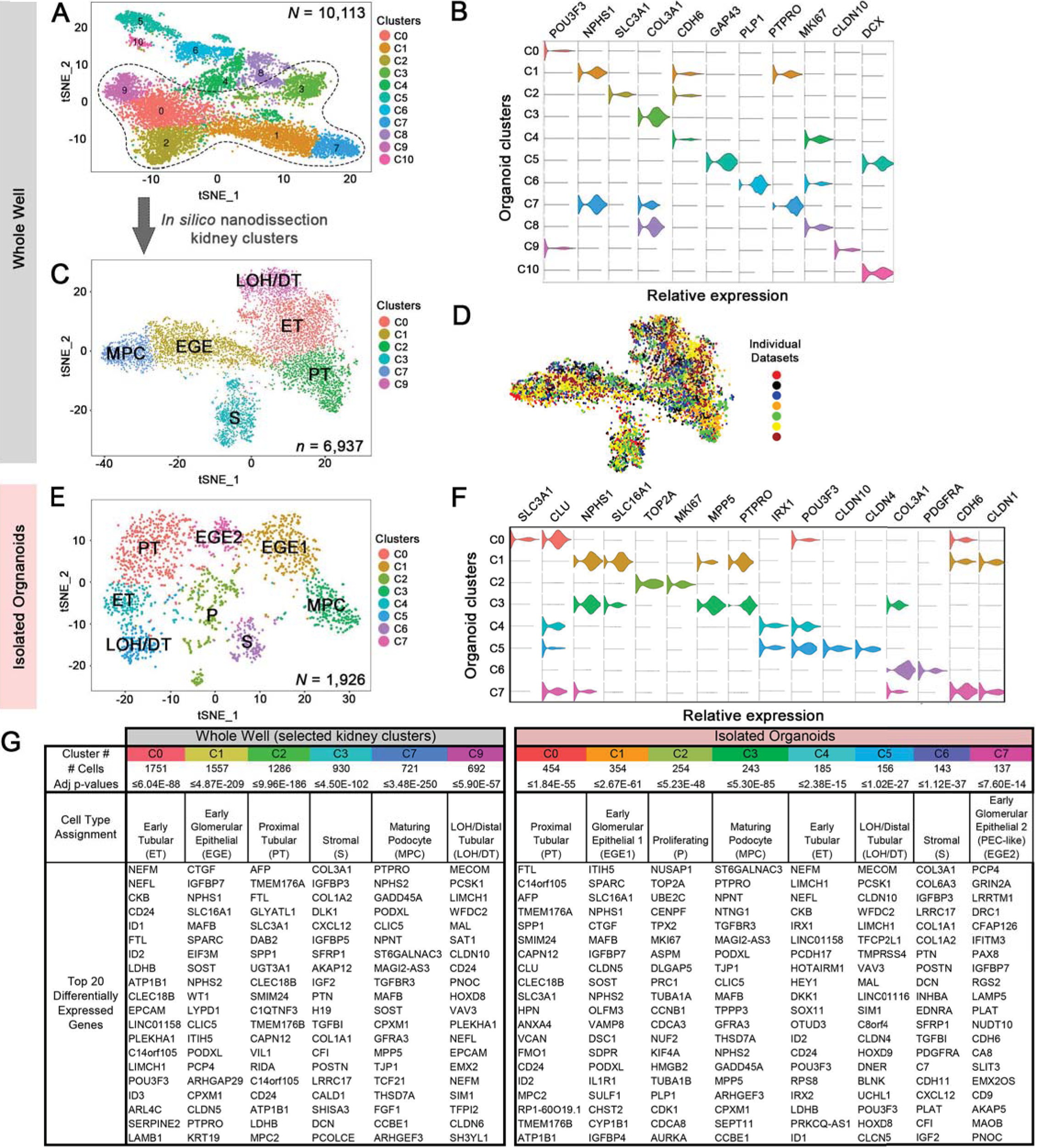
Kidney organoids profiled by single cell transcriptomics reveal spectrum of maturing nephron elements. **(A)** t-SNE plot including all cell clusters generated from scRNA-seq analysis of whole wells of kidney organoids. N, number of transcriptomes from 7 datasets from 4 individual experiments, 3 technical replicates. Dotted line indicates “kidney” clusters. **(B)** Violin plots of selected differentially expressed genes for each cell cluster for (A). **(C)** t-SNE plot showing selected kidney cell populations in whole wells of kidney organoids, identified by clustering similar single cell transcriptomes. **(D)** Overlay t-SNE plots of 7 individual datasets contributing to (C.) **(E)** t-SNE plot showing all cell populations present in isolated organoids. Data representative of 3 separate experiments. **(F)** Violin plots of genes-of-interest within cell clusters of (F). **(G)** Top differentially expressed genes within cells of these clusters compared to other cell clusters from (C) on left, and (F) on right. LOH, loop of Henle. Gene names not italicized for ease of viewing in (D, G, H). See also related Fig. S1.

Within the kidney clusters, expression of characteristic markers of podocytes (including *NPHS1*, *PTPRO and NPHS2*) was seen in two separate clusters, C1 and C7 (Fig 2B and G left). Compared to C7, relatively lower *PTPRO* and *NPHS1* as well as higher *CHD6* expression were observed in C1, suggesting that this cell cluster represents cells at an earlier stage of glomerular epithelial cell development. Thus, we designated the C1 cluster (lower *PTPRO/NPHS1*) as EGE (early glomerular epithelial) cells and C7 cluster as MPC (maturing podocyte) cells. The other clusters were consistent with ‘early’ tubule (*POU3F3*), proximal tubule (*SLC3A1*), loop of Henle/distal tubule (*CLDN10*) and stromal (*COL1A1*) cell lineages. Each of the clusters present was consistent with what we previously observed in developing human kidney (15) based on examination of the top 20 cluster-defining genes (Fig. 2B,G left).

Overlay of cells from individual transcriptional datasets revealed consistent contribution to each cell cluster by cells from each dataset (Fig. 2D). Each dataset exhibited well-dispersed cells within each cluster, confirming the reproducibility of our organoid culture system, transcriptome sequencing and bioinformatics analysis pipeline. Overall, an average of 70.3% (SD ± 9.9%) of cells from each dataset contributed to the kidney cell clusters. (Fig. S1D). Each cell-type was consistently represented, though wider variability was noted in the relative contribution to each cluster amongst datasets with SD ± 1.1% to ± 7.8% in C10 and C2 respectively (Fig. S1E). Taken together, these data indicate that organoid cultures are reproducible and include mostly kidney but also other non-kidney cell types, supporting the use of the single cell transcriptomic approach for analysis of kidney organoids to both identify and quantify kidney cell-type gene expression.

To further confirm the kidney cell-type content of kidney organoids, a second iteration of scRNA-seq was performed using isolated tubular organoids, which were microdissected from the surrounding mesenchyme (“Manually dissected Org” in Fig. 1C) (9). Unsupervised clustering of transcriptomes representing 1,926 cells revealed 8 identifiable cell clusters, similar to the “kidney clusters” shown in Fig. 2C, but did not include the “non-kidney” cell types detected in the whole well cultures with the exception of a proliferating cell cluster (Fig. 2E). A full list of genes used to define each cluster is in Table S2 while the top 20 cluster-defining genes are shown in Fig. 2G (right). (Note that a gene can be included in multiple cluster-defining gene lists since significance of gene expression by cells in one cluster is compared to expression in cells in all other clusters combined.) Quality assessment of these cells was on par with the prior combined datasets (Fig. S1F-G).

Expression of the same characteristic kidney and proliferation genes by the isolated organoid cell clusters supported the earlier assignment of cell type identity in the whole well analysis. Again, two discrete *NPHS2/NPHS1*-expressing cell clusters were present reinforcing the concept that there are two transcriptional states of podocyte-lineage cells in the organoids. The possibility that this represented two developmental stages of cells was supported by the presence of cells in either cluster expressing *CDK6*, *LHX1*, *FOXC2*, *CCNL1* and *SIX1*, indicating that these cells were still mitotically and developmentally active (19, 20).

While two *NPHS1*-expressing clusters were previously detected, EGE and MPC in whole well (C1 and C7 in Fig. 2A-B), three *NPHS1*-expressing clusters were observed in the isolated organoids (Fig. 2F, G right): one low (C7) and two high (C1 and C3). This low *NPHS1*-expressing cluster was designated as a second early glomerular epithelial (EGE2) cell cluster, and expressed markers associated with parietal epithelial cells (PECs) including *CLDN1*, *CDH6,* and *PAX8* (Fig. 2F, G right) (15). To determine whether the single EGE whole well cluster in Fig. 2C also contained such cells, subclustering was performed and revealed two clusters: one relatively high *NPHS1*/low *CLDN1* and the other relatively low *NPHS1*/high *CLDN1* expression, similar to that seen in C1 (EGE1) and C7 (EGE2) in isolated organoids (Fig. S1H-J). The many disparate cell types present in the whole well study most likely drove aggregation of these cells into a single EGE cluster despite using the same cluster resolution in both analyses (Seurat clustering parameter Res = 0.6). The lack of such non-kidney cell types in the isolated organoids allowed a more refined clustering at the same resolution. Further illustration of this difference is reflected in the variation of cluster-defining genes seen between the clustering results from whole well and isolated organoids. For example, the ET (early tubular) cluster in whole well culture shares 10 of the top 20 genes with the ET cluster in the isolated organoids. Of note is that cells with gene expression characteristic of collecting ducts, mesangial or endothelial cells were not appreciated in either cell clustering iteration. Taken together, these results support the findings of prior histologic and IF characterization of kidney organoids, reveal two discrete putative podocyte clusters and suggest that the majority of non-kidney cells reside in the mesenchyme surrounding the tubular organoids.

### Kidney organoid cells recapitulate developmental transcriptional programming of human kidney cells

To determine the similarity of gene expression in cells from hPSC-derived organoids and developing human kidney, we compared the gene expression profiles of the 6 kidney clusters from the whole well to those generated from developing human kidney (around 15 wk gestation) (21). Whole well cell clusters were chosen over the isolated organoid clusters due to the higher number of cell transcriptomes for comparison. Intriguingly, unsupervised cluster analysis of developing human kidney also revealed two distinct podocyte clusters (21), suggesting that what is observed in organoids is reflective of the developing human kidney. Indeed, averaged gene expression in MPC and EGE organoid clusters was most similar (darker red) to the corresponding clusters (Mature and Early podocyte clusters, respectively) in the developing kidney (Fig. 3A). Additionally, compared to the MPC organoid cell cluster, the EGE cluster shared greater similarity with the Proliferating cell cluster of the developing kidney, reinforcing the assertion that these cells were at an earlier developmental state.

**Figure 3.**
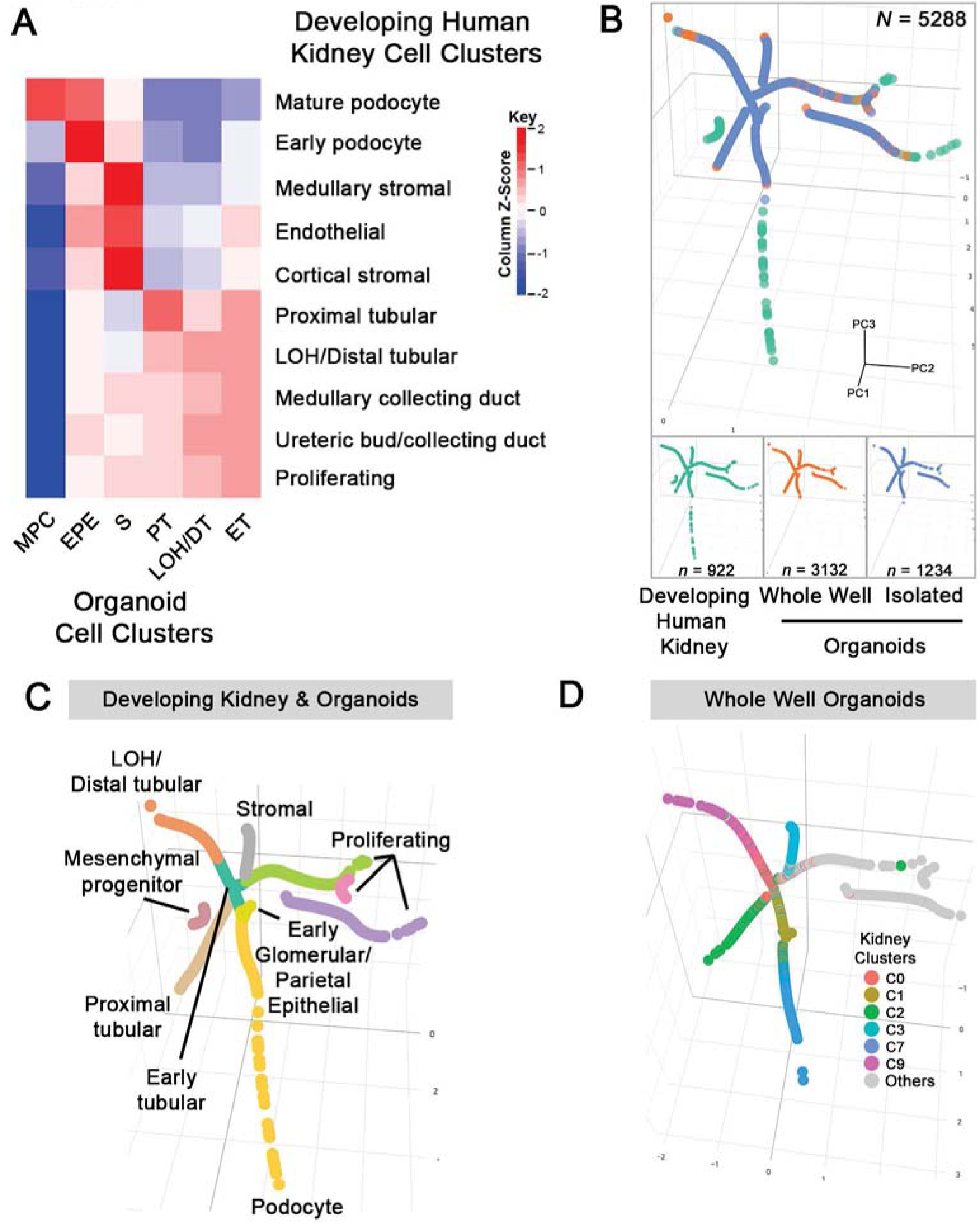
Cells of kidney organoids share the same cell-type specific transcriptional trajectories as developing human kidney. **(A)** Correlation matrix comparing gene expression profiles of cells in clusters from organoids (whole well) to developing human kidney of estimated gestation 105-115 days (27). The intensity of the overall image was adjusted to maximize the discernable visible range of Z-score intensities. **(B)** Overlaid trajectory analyses of combined single cell transcriptomes from developing human kidneys, whole well organoid cultures and isolated organoids. N, cumulative number of transcriptomes used in trajectory analysis, and *n,* contributing number of transcriptomes from each dataset. See related Video S1. **(C)** Trajectory segment cell type assignment of developing kidney and organoids based on gene expression patterns as in Fig. S2A-B. **(D)** Superimposed t-SNE cluster cell type assignments of selected kidney clusters from whole well organoids only (Fig. 2C) overlaid on cell lineage trajectory. Cells from all other clusters are represented in gray. See related Fig. S2.

Similarly, averaged gene expression of the organoid stromal cell cluster resembled stromally-derived cell clusters in the human kidney (cortical and medullary stromal, and endothelial) while averaged gene expression of tubular cell clusters of the organoids was most similar to the corresponding tubular cell clusters of the developing kidney; the early tubular organoid cell cluster shared gene expression with all tubular segments and the proliferating clusters of the developing kidney, suggesting a less differentiated cell cluster. Though gene expression correlation for tubular clusters had a lower Z-score than the previously mentioned clusters, this likely reflects variable expression of transporters and other segment-specific genes in tubular segments at different maturation stages (Table S1). Overall, the correlation analysis confirms our assignment of cell type identities to organoid kidney clusters and also demonstrates consistency between different stem cell lines (15). Moreover, this analysis further reinforces the notion that the presence of two discrete podocyte cell clusters in both organoids and developing human kidney reflects a fundamental developmental process in podocyte development.

To better define the cell types represented in organoid cultures, we sought to more directly compare cells from organoids to those from developing human kidneys. Using a second and entirely separate analysis of the single cell transcriptome data, all aforementioned organoid and developing human kidney single cell transcriptome datasets were combined and a developmental cell lineage trajectory was generated. All single cell transcriptomes were included at the outset of this analysis, but the trajectory algorithm dispensed with cell types not well represented in both organoids and developing human kidneys. This approach allowed the examination of cell lineages of organoids within the context of cells from developing kidneys, and of contribution of cells found in isolated organoids relative to the whole well organoid cultures (Fig. 3B, Video S1).

Contribution to each cell lineage trajectory was seen by both developing kidney and organoids. Top differentially expressed genes in each trajectory segment served as the basis for cell lineage assignment (Fig. S2A). As shown in Fig. 3C, multiple cell lineages radiated out from a central hub of precursor cells, including podocyte, proximal and distal tubular, and stromal as well as non-contiguous proliferating cell lineages in a parallel fashion between human developing kidney and organoids. The podocyte lineage further diverged into two separate lineages, maturing podocytes and PECs, which were distinguishable by their different expression profiles of four characteristic PEC genes (*CLDN1*, *ANXA3*, *PAX8* and *CDH6*) (Fig. S2B). Cells from developing kidney cells (green in Fig. 3B) contributed to the distal ends of each of the radiating cell lineage trajectories, indicating that developing kidneys contain cells that are relatively more mature than those found in organoids of this vintage.

To determine the inter-reliability of the two independent analytic methods of the transcriptomic data, cluster-assigned cells were overlaid onto the combined trajectory. We specifically used the kidney clusters from the whole well organoid clusters for this analysis to further assess the veracity of our earlier cell type assignments to each cluster, and to further explore the basis of the separation of the *NPHS1/NPHS-2*-expressing cell clusters. Cluster-assigned cells of the kidney organoid clusters were overlaid onto the trajectory analysis from Fig. 3C, and shown in Fig. 3D. Importantly, cell cluster assignments correlated reliably with cell lineage trajectory. Cells from the EGE cluster (C1, gold) populated the early segment of the combined PEC/podocyte trajectory, while those in the MPC cluster localized solely to the post-PEC divergence segment (C7, blue). This observation indicated that cells in the EGE cluster included precursor cells common to both podocyte and PEC lineages, which suggested a basis for the segregation of EGE and MPC cell clusters. Moreover, the early overlap and proximity of the PEC and podocyte trajectories reinforced the concept that these cell types share significant transcriptional programming during development. The gene expression patterns revealed in these trajectories persist into adulthood. Indeed, expression of *WT1* and *PTPRO* expression in PEC and podocyte lineages (Fig. S2C) was reflected on a protein expression level in both PECs (WT1+/PTPRO- cells lining Bowman’s capsule) and podocytes (intraglomerular WT1+/PTPRO+ cells) in adult human kidney (Fig. S2D).

The segmentation of early and later developmental stages seen in podocytes was repeated in tubular cell lineage trajectories (Fig. 3D). Cells from the Early Tubular cluster (C0, salmon) localized more centrally, while those from the Proximal Tubular (C2, green) and LOH/DT (C9, purple) clusters localized more peripherally. To determine which organoid cells the algorithm included in the trajectory analysis, cells were mapped back onto their corresponding tSNE plots (Fig. S2E). This revealed that cells from each cell cluster contributed to the trajectory, with the non-kidney clusters contributing to the proliferating lineages. Taken together, these data indicate that cells in kidney organoids reliably recapitulate the developmental transcriptional programming observed in homologous cell types of the developing human kidney.

### Organoid podocyte cell clusters demonstrate distinct transcriptional states

We next sought to further characterize the transcriptional program in the two podocyte clusters to understand the nature of their segregation. The EGE and MPC clusters together represented 22.5% of all cells in organoid cultures (Fig. 2C, S1E) and both were characterized by expression of typical podocyte genes, including *PTPRO*, *NPHS1*, *NPHS2*, *PODXL*, and *WT1* (Fig. 4A). These two cell clusters differed, however, by the relative expression of epithelial polarity genes *MPP5*, *TJP1*, *PARD3B*, *IQGAP2*, with MPC exhibiting higher expression than EGE indicating that MPC cells were more polarized and thus at a later stage of maturation. This was confirmed on a protein expression level with the tight junction gene ZO-1 (*TJP1*), which was found only in a subset of nephrin+ cells in organoids (Fig. 4B). When these genes were examined in the context of podocyte-lineage trajectory (from Fig. 3C), expression of these genes was noted to increase in sync with the developmental trajectory, with podocyte marker genes preceding polarity gene expression (compare top and bottom rows in Fig. 4C).

**Figure 4.**
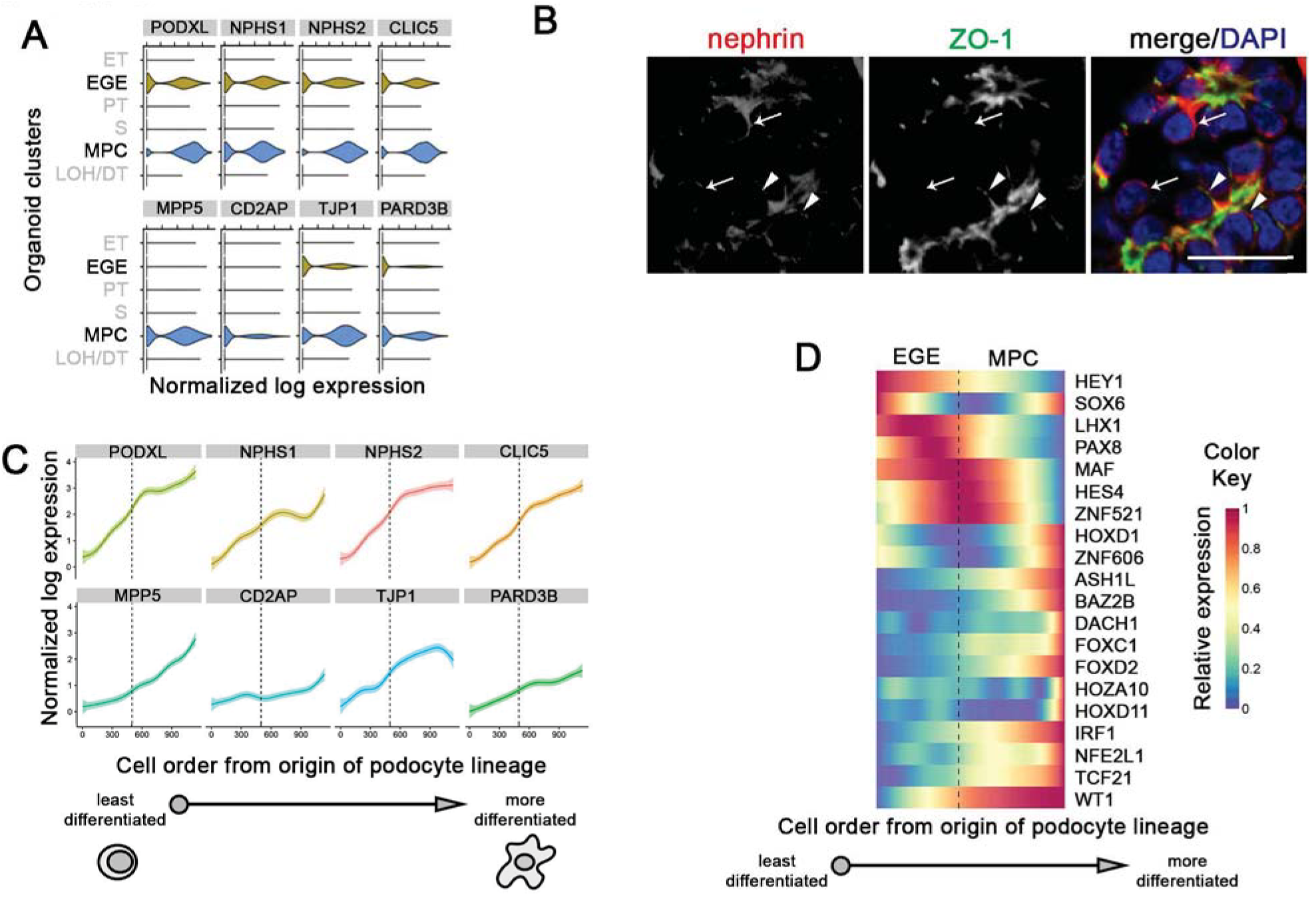
Cells farther along podocyte lineage trajectory are more mature. **(A)** Violin plots depicting expression of quintessential (top row) podocyte and (bottom row) epithelial cell polarity genes in organoid clusters from Fig. 2C. Ticks on x-axis indicate whole integers starting from 0 on the left of each plot. **(B)** Immunofluorescence confocal images showing protein expression in nascent podocytes in 20 day organoids. Arrows highlight nephrin^+^/ZO-1^-^cells while arrowheads highlight nephrin^+^/ZO-1^+^ cells. ZO-1 *(TJP1),* bar = 20 μm **(C)** Average expression of indicated genes by cells along combined organoid-developing kidney podocyte lineage trajectory from Fig. 3C using a curve fitting method (GAM) with 95% CI represented by shading. **(D)** Heatmap of gene expression of transcription factors along the podocyte lineage trajectory as in (C). Vertical dotted line indicates position at which podocyte and PEC lineages diverge (C and D). See related Fig. S2.

To determine if transcription factors (TFs) were contributing to the transcriptional segregation of podocyte clusters, the expression pattern of all known 1,639 human TFs (22) was assessed in cells along the podocyte trajectory (Fig. 4D). A subset (7 of 20 total TFs expressed in the podocyte lineage) of these was included in the differentially expressed genes in EGE and MPC clusters. While regulation of gene expression is specific tissues is much more complex than a reflection of expression levels of TFs (23), co-expression of transcription factors and their targets is frequently observed (24, 25) and at least consistent with potential regulatory patterns present. Consistent with established developmental patterns, *LHX1* and *PAX8* expression peaked earlier in podocyte development, while expression of several TFs described as involved in podocyte maturation was seen later, including *FOXC1* and *TCF21* (26, 27). An increase in expression of *WT1*, *HES4* and *MAF* around the divergence of the podocyte and PEC lineages suggested a possible basis for a regulatory transcriptional “switch” associated with podocyte maturation. Together, the trajectory analysis and gene expression characterization indicate that the EGE and MPC cell clusters represent two transcriptionally discrete states within the continuum of podocyte development.

### Genes highly expressed in immature glomerular epithelial cells of organoids are dysregulated in human kidney disease

We hypothesized that the gene expression pattern seen in the EGE cluster is reactivated in injured podocytes in glomerular disease. To test this hypothesis, genes unique to or shared by both podocyte lineage clusters (C1 and C7 in Table S1) were identified. This resulted in three sets of genes: EGE (69 genes), Shared (104 genes), and MPC (168 genes) (Fig. 5A). These gene sets were used to generate “aggregate gene expression scores” in isolated glomerular tissue from a cohort of individuals with various etiologies of chronic kidney disease (European Renal cDNA Bank, ERCB)(28, 29). Query of the subset of the identified genes available on the microarray platforms (Fig. 5A) revealed that the EGE aggregate score was significantly increased in the 170 individuals with chronic kidney disease relative to kidneys from living donors, specifically in individuals with lupus nephritis (LN), diabetic kidney disease (DKD), ANCA-associated vasculitis (AAV) and focal segmental glomerulosclerosis (FSGS) (Fig. 5B, S3A). Conversely, the MPC and Shared aggregate scores were unchanged. These results suggested that the EGE aggregate score was capturing a pathologically relevant transcriptional state in human glomerular disease.

**Figure 5.**
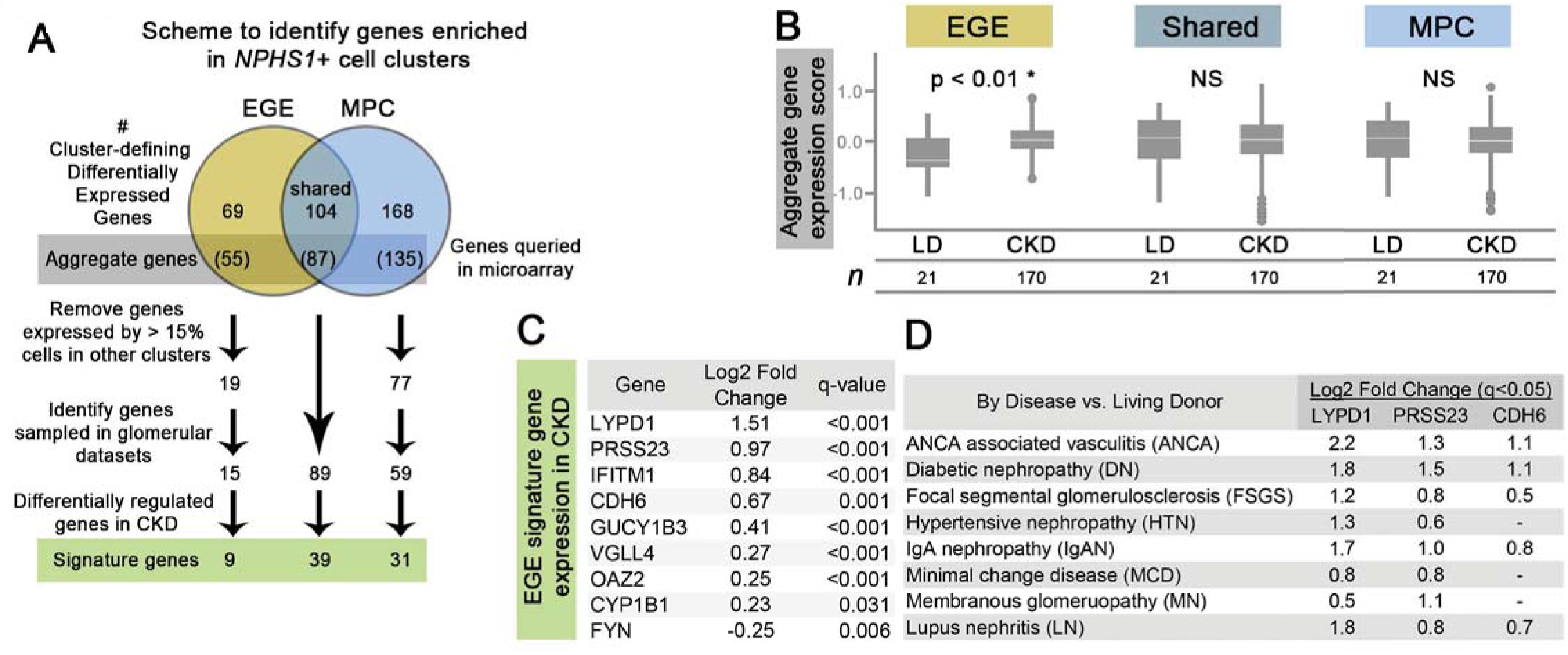
Genes differentially expressed in podocytes in organoids correlate with human kidney disease. **(A)** Cartoon depicting method of identification of genes relatively specific to early glomerular epithelial (EGE) and maturing podocyte (MPC) dusters from Fig. 2C. **(B)** Aggregate gene expression scores for three sets of genes from A (gray box) in microdissected glomerular tissue from individuals with chronic kidney disease (CKD) compared to living donors (LD) from the ERCB cohort. *n,* number of individuals. Statistical significance was calculated using two-tailed student’s t-test. Box plots indicate 75th percentile, white lines indicate median expression, asterisk denotes p < 0.05. **(C)** List of EGE “signature genes” from A that are significantly differentially expressed in CKD compared to LD. Differential expression statistics were calculated using SAM, q-values shown. **(D)** Expression of three genes from C reported by specific subsets of CKD. Gene names not italicized for ease of visualization. See also Fig. S3.

To identify potential mechanistic pathways associated with these gene sets, pathway analysis was performed. Canonical pathways generated using all of the 143 EGE genes modestly favored inhibition of angiogenesis by TSP1, hypoxia signaling in the cardiovascular system and ephrin receptor signaling, while those generated using all of the 272 MPC genes more evidently favored unfolded protein response, axonal guidance signaling and tight junction signaling (Fig. S3B). Only 16 genes of the “unique” 69 EGE subset were found in any known pathway, indicating that these genes are not well represented in known biological networks (Fig. S3C, top). This improved only modestly by expanding to include the full 143 EGE gene set (Fig. S3C, bottom). Thus, pathway analysis of EGE genes provided limited information about potential pathogenic mechanisms of disease, pointing to the need to examine the contribution of individual genes in the EGE set.

A gene prioritization approach was used to identify candidate genes driving the EGE signature expression in CKD. Genes enriched in the EGE cluster were identified when expression in all other clusters combined was below 15%, indicating that expression was relatively unique to the EGE cluster cells. The same threshold was used to identify MPC-enriched and Shared-enriched genes (Fig. 5A). Application of this algorithm identified 19 genes whose expression was enriched in the EGE cluster, 15 of which were present and analyzed on the microarray platforms used to interrogate the transcriptomes of human kidney glomerular tissues. Of the 15, expression of 8 was significantly increased while 1 was decreased in CKD (Fig. 5C). Meanwhile, 31 of 59 MPC-enriched genes and 33 of 89 Shared-enriched genes were differentially regulated in CKD (Fig. S3D). Notably, expression of *MPP5*, *IQGAP2*, *NPHS1*, *PTPRO*, *TJP1* and *WT1* was decreased in CKD.

Examination of the EGE gene signature (Fig. 5C) revealed that the function of several gene products could affect the course of glomerular disease, although detailed literature especially pertaining to the functions of these proteins within the kidney are lacking (Table S3). A unifying theme for these gene functions is development and cell survival. Thus, by employing a subtractive approach, a small set of EGE genes was identified, the majority of which associated with significantly increased gene expression in diseased glomeruli. Together with the finding that expression of genes characteristic of mature podocytes were decreased, these data suggest podocyte loss or dedifferentiation was associated with expression of EGE signature genes.

### Increased expression of *LYPD1*, *PRSS23* and *CDH6* correlates with human glomerular disease

We focused on three significantly differentially regulated genes of the EGE list associated with CKD. The first two (*LYPD1* and *PRSS23*) showed the most significant differential expression in disease, while the third (*CDH6*) has a well-described role in kidney development and was therefore of particular interest (30). Expression of each was variably increased in subgroups of individuals with different etiologies of CKD (Fig. 5D). As expected, cells expressing these genes in organoids were predominantly located in the EGE cluster (Fig. 6A). The pattern of expression of these genes was also examined in the context of the podocyte-PEC lineage trajectory from Fig 3C and revealed that expression of each of these genes was decreased in maturing podocytes, but was increased in common precursor and PEC-lineage cells (Fig. 6B). These results indicate that the combination of increased gene expression of *LYPD1*, *PRSS23,* and *CDH6* is consistent with a less differentiated podocyte or PEC.

**Figure 6.**
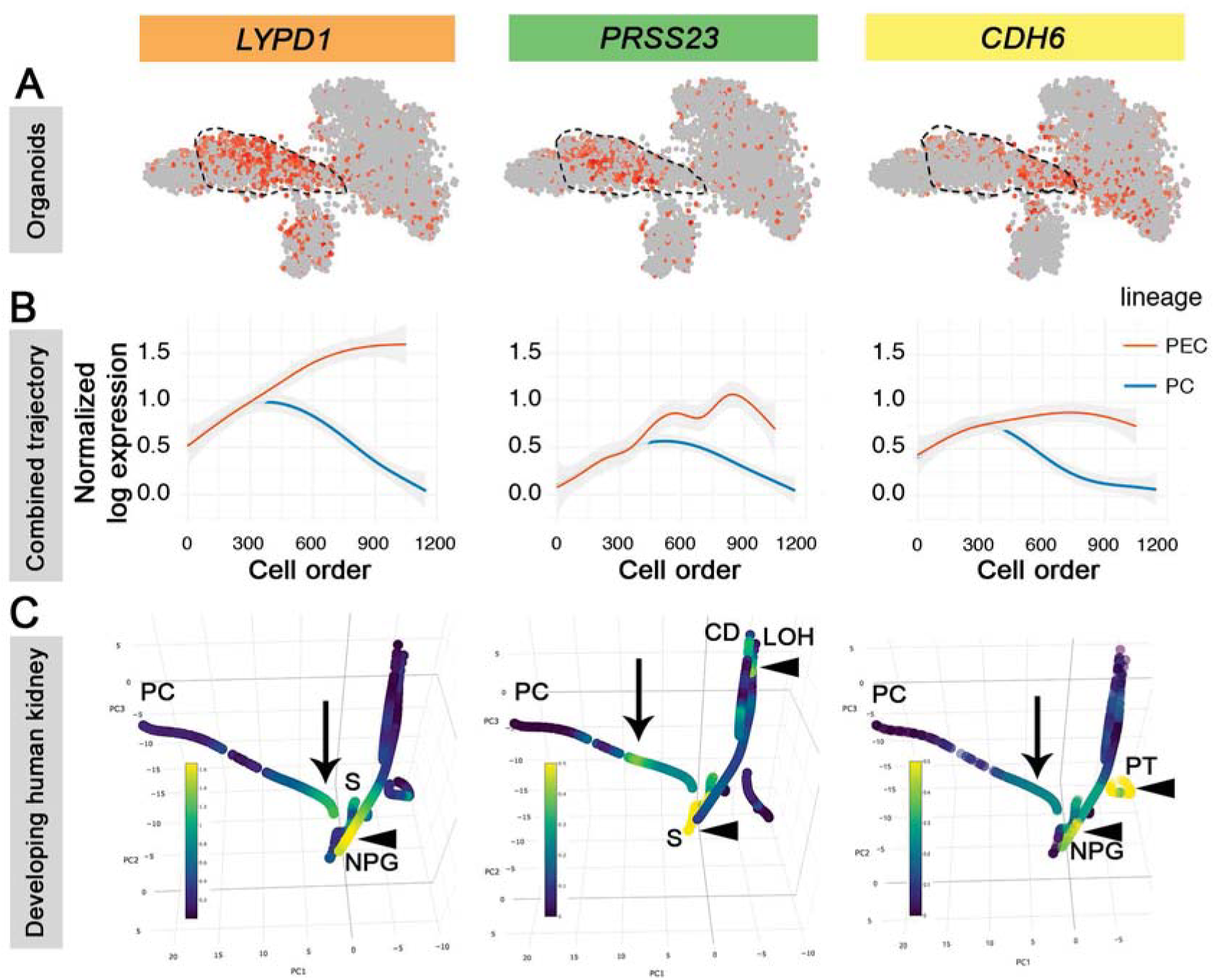
Expression of *LYPD1, PRSS23,* and *CDH6* declines during podocyte maturation in organoids and human kidneys.

**(A)** Feature plots of *LYPD1* (left), *PRSS23* (middle) and *CDH6* (right) expression overlaying kidney organoid tSNE plot from Fig. 2C. Dashed lines outline cells in EGE cluster. **(B)** Average expression of *LYPD1* (left), *PRSS23* (middle) and *CDH6* (right) in cells of overlaid podocyte (*n* = 645 transcriptomes) and PEC (*n* = 546 transcriptomes) lineage trajectories, color key on far right. Gray shading represents 95% CI. **(C)** Developing human kidney trajectories as described in Menon et al. (31) representing 6,414 transcriptomes with overlaid gene expression of *LYPD1* (left), *PRSS23* (middle) and *CDH6* (right) oriented for optimal visualization of podocyte lineage, arrows highlighting maximal podocyte lineage expression, arrowheads highlighting non-podocyte lineage maximal expression. Cell lineage labels: PC, podocyte; S, stromal; NPG, nephron progenitor; CD, collecting duct; LOH, Loop of Henle; PT, proximal tubule. Color scales represent normalized log expression.

To explore why these three genes were not identified as enriched in early podocyte-lineage cells in our earlier study of single cell transcriptional profiling of the developing human kidney (31), we specifically interrogated gene expression in the context of the developing kidney cell lineage trajectory analysis generated in that study. Examination of the three gene expression patterns revealed that each gene was expressed by cells early in podocyte lineage as well as other cell type lineages: *LYPD1* expression was found in nephron progenitor (NPG) cells, *PRSS23* expression was found in stromal, CD and LOH lineage cells, and *CDH6* was found in PT and NPG lineage cells (Fig. 6C). Thus, these three genes were indeed expressed in developing podocytes in human kidney, but expression in other cell types masked their recognition. So the podocyte-enriched organoid cultures revealed insights about podocyte development that were not apparent by solely examining the list of differentially-expressed genes in the cell clusters of the developing human kidney (31).

Expression of *LYPD1*, *PRSS23* and *CDH6* was investigated in diseased kidney tissue. Reiterating Fig. 4C, expression of each was increased in various specific subgroups of individuals with CKD from the ERCB cohort, here shown more descriptively by box plot (Fig. 7A). More specifically, expression of each also correlated positively with markers of CKD progression including loss of eGFR (Fig. 7B) and presence of nephrotic range proteinuria in lupus nephritis (Fig. 7C). In contrast, expression of genes from the MPC (*MPP5*, *IQGAP2*) and Shared (*NPHS1*, *WT1*, *PTPRO*, *TJP1*) sets correlated positively with eGFR (Fig. S4A). These results are especially poignant as this means that increased expression of the three signature genes accompanied loss of kidney function and loss of characteristic podocyte marker expression.

**Figure 7.**
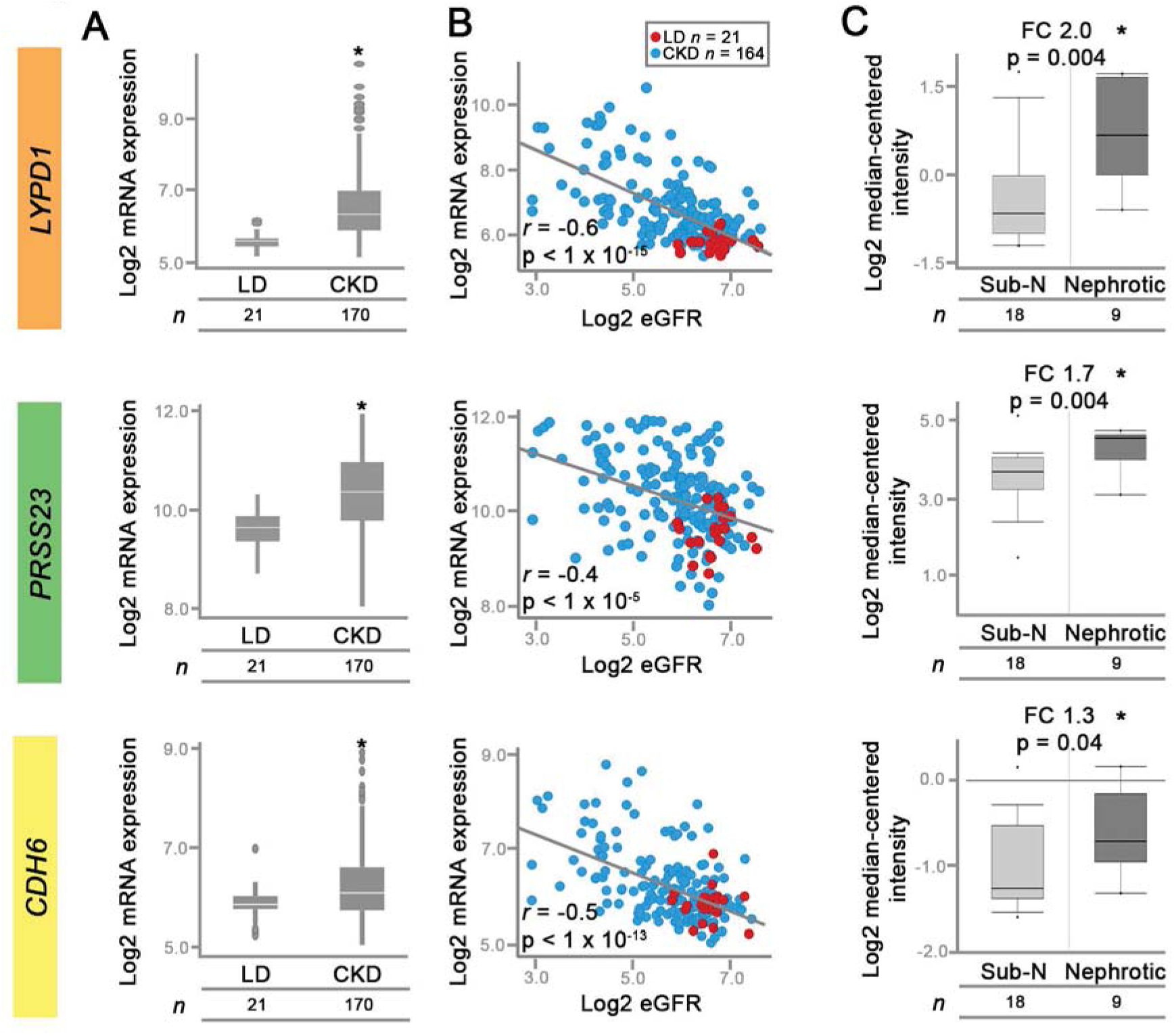
Expression of candidate early glomerular epithelial/PEC genes correlates with chronic kidney disease. **(A)** Box plots comparing *LYPD1* (top), *PRSS23* (middle) and *CDH6* (bottom) expression in CKD relative to LD, as in Fig. 5C. **(B)** Scatter plots showing gene expression from (A) relative to eGFR in CKD and LD. Statistical significance calculated using Pearson correlation, associated p-values shown. **(C)** Box plots comparing individual gene expression levels to proteinuria groups in lupus nephritis samples from ERCB in Nephroseq (www.nephroseq.org). Statistical significance calculated using one-tailed t-test, associated p-values shown. Sub-N, sub-nephrotic; FC, fold change; *n,* number of individuals. See also Fig. S4.

To validate these findings, expression of these three genes was also examined in a separate cohort of individuals with glomerulonephritis including nephrotic syndrome from NEPTUNE (*n* = 90) and antineutrophil cytoplasmic antibody-associated vasculitis (*n* = 15) (28). *LYPD1*, *PRSS23* and *CDH6* expression again correlated inversely with eGFR at baseline (acquired within 4 weeks of kidney biopsy) (Fig. S4B) but only *PRSS23* expression significantly correlated with proteinuria in the nephrotic individuals for whom UPCRs were available (Fig. S4C). However, when the nephrotic group was analyzed by etiology, all 3 genes significantly correlated with proteinuria in the membranous nephropathy subgroup (n = 44) (Fig. S4D). Additionally, expression of *LYPD1*, *PRSS23* and *CDH6* was examined in isolated glomerular tissue from a rat model of focal segmental glomerulosclerosis (FSGS) (Fig. S4E) (32). Here, gene expression levels are reported relative to podocin (*Nphs2*) and nephrin (*Nphs1*) expression levels as performed previously. Again, all three genes showed increased expression relative to podocyte genes in the FSGS rat as compared to the control rat tissues. Thus, both a second cohort of podocytopathies and a rat model of FSGS also showed that expression of these three signature genes was increased in glomerular disease.

To address which cell type within the dissected glomeruli may be contributing to the increase in expression of these genes, *PRSS23* expression was examined in diseased human kidney tissue (Fig. 8), as it appeared to have the most robust association with both proteinuria and renal function in our earlier studies. ISH revealed that *PRSS23* transcripts were detected in cells lying along Bowman’s capsule in both control and diseased kidney tissue consistent with parietal epithelial cells (Fig. 8, arrowheads). However, in diseased kidney, *PRSS23* transcripts were also detected in cells within the glomerulus present along the urinary surface (Fig. 8, arrows).

**Figure 8.**
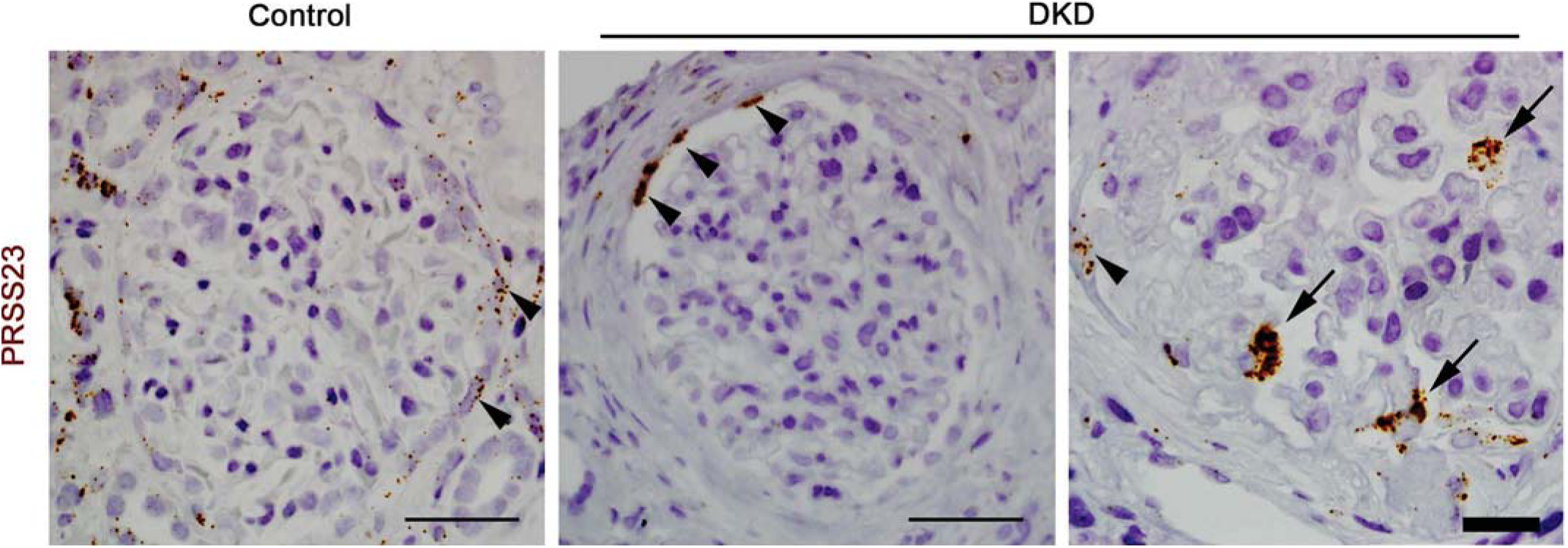
Expression of PRSS23 in diseased human glomeruli. *PRSS23* expression as detected by ISH in control and diabetic kidney disease (DKD) sections, arrowheads highlight staining of cells at the glomerular parietal surface, arrows highlight cells on the visceral surface, thin black bar = 50 μm, thick black bar = 20 μm. Representative images of 5-10 glomeruli from each sample, and 2 separate DKD samples.

Our results show that a gene expression signature defined in early glomerular epithelial cells in kidney organoid cultures was detected in human glomerular disease. The gene signature included *LYPD1*, *PRSS23* and *CDH6*, each of whose expression was increased in a cohort of glomerular disease, findings which were recapitulated in a second human glomerular disease cohort, as well as a rat model of FSGS. Taken together, expression of these genes in human glomerular disease is consistent with an activation of a transcriptional program also seen in developing glomerular epithelial cells in organoids and human kidney.

## DISCUSSION

Podocyte injury and adaptation are central to progressive decline in kidney function in glomerular disease but identifying the molecular events underlying this process in human disease has been challenging. We describe an early glomerular epithelial cell gene expression signature generated from single cell transcriptional profiling of human PSC-derived kidney organoids, and show that this gene set is differentially expressed in diseased human glomeruli. By combining cell clustering and cell lineage trajectory analyses of organoid and developing human kidney single cell transcriptomes, we showed that cells in two distinct *NPHS1/NPHS2-*expressing organoid clusters shared the same developmental transcriptional program with human podocytes *in vivo* but represented different stages of podocyte development, revealing that the gene signature arose from early glomerular epithelial cells. Identification of a glomerular-disease relevant gene expression signature corresponding to the early developmental podocyte stages supports the concept of transcriptional plasticity as a compensatory mechanism in glomerular disease.

Most of the nine early glomerular epithelial signature genes were not previously appreciated as being involved in glomerular pathology and several are minimally annotated, revealing new gene targets for investigation. Of the three signature genes (*LYPD1*, *PRSS23* and *CDH6*) that were found to be associated with proteinuria and loss of kidney function, *PRSS23* is of particular interest given this serine protease’s proposed role in Snail-dependent epithelial to mesenchymal transition (33). We found that *PRSS23* expression was increased in developing EGEs and PECs relative to podocytes, but only *Cdh6* (of the nine signature genes) was increased in healthy PEC-enriched rat glomerular isolates (34). This suggests that expression of these genes in our diseased human glomerular tissues does not simply represent inclusion of healthy PECs in the glomerular tissue samples. In DKD glomeruli, cells expressing *PRSS23* were seen both at Bowman’s capsule and the urinary glomerular basement membrane surface. The influx of cells from Bowman’s capsule is consistent with prior descriptions of podocytopathies in humans (35-37) and was shown to be of PEC origin by marker expression in human tissue and lineage tracing in a mouse FSGS model (38). An independent mouse model of FSGS showed that *Prss23* expression was significantly increased in lineage-tagged podocytes isolated six weeks after disease induction (39), though it is possible that expression may be due to trans-differentiating PECs that express *COL1A1*, which was used in the lineage tag. Together, these observations support the intriguing possibility that *PRSS23* is involved in the development of a migratory PEC-podocyte phenotype in glomerular disease. Other possible cells responsible for the signature gene expression include trans-differentiating renin-lineage cells (40), which can migrate across the glomerular or mesangial basement membrane. However, our description of these nine genes within the context of developing podocytes provides more compelling evidence in support of PEC-podocyte phenotype plasticity.

In addition to identifying a novel disease signature, our comparative trajectory analysis fills a critical knowledge gap by validating that cells in human PSC-derived organoids faithfully reproduce the genetic developmental program of podocytes as well as other cell types in kidney organoids. The organoids recapitulated gene expression in cells of developing human kidneys, and led to the description of a gene expression signature unique to developing podocytes. Surprisingly, though developing podocytes expressed these genes *in vivo*, some were identified only by differential transcriptional profiling in developing podocytes isolated from kidney organoids as opposed to developing kidneys themselves, due to the more targeted analysis in organoid cultures. Our analysis also revealed the complexity of molecular signatures distinguishing podocyte dedifferentiation versus PEC trans-differentiation as illustrated by our trajectory analysis in which podocyte and PEC developmental lineages emerged from a common glomerular epithelial precursor (41). Divergence of these two cell lineages may be due to a transition of transcriptional programming similar to what was recently shown for Notch signaling driving intercalated to principal cell transition (42), a concept which is supported in our current study by the alterations in transcription factor expression patterning along the podocyte trajectory. Comprehensive analysis of transcription factor patterning in podocyte-PEC development may help discern these lineages more fully, and when this knowledge is applied to glomerular disease, may aid in identification of upstream regulators of disease-driving transcriptional activity.

Our dual transcriptional characterization of the repertoire of cells generated in PSC-derived kidney organoids also elucidates how each analytic method contributes complementary insights regarding kidney organoid cultures as well as kidney development and disease. While the trajectory analysis confirmed cell types and vintages of the organoid cells relative to developing human kidney, cell clustering analysis highlighted cell-type specific gene expression profiles as well as provided intriguing clues regarding cell-lineage development, e.g. two separate clusters of podocytes corresponded to different segments of podocyte development as discussed above. Cluster analysis also revealed kidney cells (including podocytes, tubular, and stromal) could be highly enriched by separating the tubular organoids from the surrounding mesenchyme, whereas non-kidney cell types identified in previous characterizations (15, 43, 44) were successfully depleted. However, both single cell transcriptional analyses showed that kidney cell types generated in organoid cultures were reproducible and robust. Thus, our study demonstrates that single cell transcriptional characterization of kidney organoids can be exploited in multiple ways, including defining and refining the cellular complexity of organoids and discovering transcriptional programmatic themes.

Although this work represents a significant advance in our understanding of kidney organoid transcriptional states, several factors limit interpretation of our results and warrant future investigation. Mesangial and endothelial cells were not described in this analysis, potentially as a result of developmental immaturity. Conversely, several cell types not of kidney origin were found. Both observations are consistent with prior results (15, 43). However, we showed that non-kidney cells could be limited by either manually isolating organoids from the surrounding mesenchyme or by post hoc *in silico* nanodissection of cells-of-interest. An additional observation was that organoid kidney cells corresponded to earlier developmental stages of human developing kidneys in late first trimester to mid second trimester (not surprising for cells differentiated less than 3 weeks in culture), meaning late stages of nephron development are not well represented. Indeed, prior studies showed that PSC-derived organoid podocytes are similar to mammalian podocytes at the capillary loop stage based on cytoskeletal architecture and marker expression (17). Together, identification of such factors provides a framework to direct further optimization of organoid differentiation protocols.

In conclusion, in addition to expanding our knowledge of transcriptional events during podocyte development, our approach has unveiled several novel glomerular disease-associated transcriptional programs. Further exploration regarding the basis of this developmental glomerular gene signature may aid in identification of novel glomerular disease biomarkers and treatment strategies. Deciphering such events in diseased human glomeruli, often accompanied by manifestations of advanced disease such as sclerosis and fibrosis, is challenging. In contrast, human kidney organoids are a simpler system incorporating key glomerular cell types such as podocytes and PEC, which are highly amenable to purification, treatment with small molecules, and omics-scale analysis. Moreover, characterizing kidney organoids on a single cell transcriptomic level provides *a priori* knowledge of gene sets available for experimental manipulation *ex vivo*. Thus, a previously unrecognized beneficial role of PSC-derived kidney organoid cultures to identify novel biomarkers of glomerular specific kidney diseases is revealed. Our results highlight the utility of kidney organoids as a discovery tool to define and investigate pathomechanisms of glomerular disease.

## METHODS

### Generation of kidney organoids

Kidney organoids were generated using UM77-2 hESCs (NIH Registration# 0278) as described (9, 15). Mycoplasma contamination-free status of actively passaged cell lines was confirmed prior to differentiation using the Universal Mycoplasma Detection Kit (ATCC, Manassas, VA, 30-1012K) per manufacturer’s protocol. Organoids for immunocytochemistry were picked 20-27 d post-plating, fixed in 4% paraformaldehyde (Electron Microscopy Sciences, Hatfield, PA)/PBS (Gibco), infiltrated with a sequential gradient of sucrose in PBS and embedded in 20% sucrose/OCT (Tissue-Plus, Fisher HealthCare) as previously described (49).

### Human kidney tissues

Human control kidney tissue (tumor nephrectomy) was procured and prepared via the tissue procurement service at the University of Michigan Comprehensive Cancer Center. Normal human kidney tissues were obtained from protocol pre-transplant donor biopsy, and human diabetic kidney tissues were obtained from individuals then procured by, routinely processed by and accessed through the archives of the Dept. of Pathology at the University of Michigan. All tissues were formalin fixed, paraffin embedded and sectioned at 3 μm thickness.

### Immunostaining and imaging of organoids and human kidney tissues

Cryosections were rehydrated with PBS, blocked with 5% normal donkey serum (Jackson Laboratories) plus 0.1% Triton X-100 (IBI Scientific, Peosta, IA) in PBS, and immunostained with antibodies in 3% BSA (Cohn Fraction V, Sigma-Aldrich, St. Louis, MO) in PBS. Sequential WT1 and PTPRO (Glepp1) staining of the same section of normal human kidney (donor) protocol biopsy was performed as previously described except that anti-WT1 was substituted for anti-TLE4 antibody to identify podocyte nuclei (50, 51). Species appropriate fluorescence-tagged or peroxidase conjugated secondary antibodies were used to detect primary antibodies. Samples were mounted using SlowFade or Prolong Gold with DAPI (Invitrogen). DAB was used to develop peroxidase product and counterstained with hematoxylin. Images were acquired using indicated light, epifluorescence or confocal microscope with digital camera.

### ISH of human kidney tissue

RNAscope probe for Hs-PRSS23 was obtained from Advanced Cell Diagnostics (Newark, CA, catalog #506571) and samples were prepared, probed and counterstained per the manufacturer’s protocol.

### Rat FSGS model tissue and glomerular transcriptional data

Frozen kidney tissue from sham nephrectomized wild type and unilaterally nephrectomized TG Fischer344 rats (expressing the AA-4E-BP1 transgene under the control of the human podocin promoter) was previously generated as described (32) and cryosectioned. Microarray data were previously generated as part of the same study from isolated glomeruli using Rat Gene ST 2.1 Affymetrix gene array and reanalyzed for genes of interest using the described method (32).

### Antibodies

Alexa-FLUOR labeled secondary Abs were obtained from Invitrogen. Peroxidase-conjugated mouse IgG kit was obtained from Vectorstain (PK-6102). Primary antibodies were sourced as follows: Nephrin (R&D, AF4269); Cadherin-6 (R&D, MAB2715); MEIS1/2/3 (Active Motif, ATM39795; Podocalyxin (R&D, AF1658), WT-1 (Abcam, ab89901); PTPRO (5C11, gift of R.C.W. (52)); N-cadherin (R&D, AF6426); ZO-1 (Invitrogen, 40-2300); Pan-cytokeratin (Sigma, C2562); E-cadherin (BD, 610181); PRSS23 (Sigma, HPA030591).

### Organoid single cell RNA sequencing and bioinformatic analysis

Kidney organoids were harvested between 18-21 D in culture, as either whole well or isolated spheroids (Fig. 1B). Cells were dissociated using cold active protease into single cells and Drop-seq was performed as previously described (15). Single cell transcriptomes from 7 datasets generated from 4 separate kidney organoid culture experiments (3 biological replicates) were included in the “whole well” analysis, while a single dataset was included in the “isolated organoids” analysis. Organoid scRNA-seq data matrix preprocessing, normalization, log transformation, unsupervised cell clustering, scaling, PCA dimensionality reduction, highly variable gene identification, individual gene expression query and presentation, and correlation with cell clusters previously derived from developing human kidney scRNA-seq (31) were all performed using the Seurat R package as described (15, 31, 53). Cells expressing under 500 or over 4000 genes or above 25% mitochondrial reads were excluded in the analysis. Seurat ScaleData function was used to regress out technical variables including mitochondrial read content, number of UMI per cell and batch effect. Unsupervised clustering was performed with a resolution level of 0.6. tSNE clustering was used for wo-dimensional frame viewing of clusters (54). Default non-parametric Wilcoxon rank sum test was used for differential expression analysis.

### Combined developmental trajectory cell lineage from single cell transcriptomes

Trajectory analysis was performed with a method developed by Zhou and Troyanskaya (31). Cells expressing at least 1000 genes were included for analysis. To control for technical variations within the single cell transcriptomic data, linear modeling in Seurat was used to regress out correlation with the number of genes expressed, percentage of mitochondria reads, and batch variables. To control for confounding effects from cell cycle, the cell cycle signal was regressed out using regularized linear models as previously described (31). Top principal components were used for trajectory analysis. Joint trajectory analysis combined human developing kidney (GEO accession GSE109205) and organoid datasets (7 “whole well” plus 1 “isolated organoid”) as follows. Each was individually processed and technical variations removed as described above, then all were aligned using the canonical correlation analysis (CCA) method as implemented in Seurat; the canonical component projections were used instead of the principal component projections for the joint trajectory analysis. Developing human kidney and organoid data were weighted equally by the algorithm, resulting in agnostic exclusion of cell types not well represented in both data sources. Subsequent gene expression plots were generated from the trajectory analysis as follows. Heatmaps were generated from genes with average log-scaled expression > 0.1 and p-values of differential expression along the trajectory < 0.001 (two-sided Wald test). Scaled log expression was generated by generalized additive modeling (GAM). Individual gene expression plots were generated from average cell gene expression along the trajectory, log values of which were normalized, using a curve fitting method (GAM) with 95% CI. For data presentation referring to “cell order,” cells are aligned sequentially based on the pseudotime order from the estimated trajectory.

### Transcriptomic analysis of human kidney tissue

The discovery cohort included previously generated microarray data from microdissected human glomeruli and sourced from individuals with kidney disease (n=170) and healthy donors (n=21) from ERCB (55) which were and accessed in GEO by accession GSE104948. The validation cohort (n=111) also included previously generated microarray data from microdissected glomeruli from individuals with FSGS (n=30), MCD (n=15), MN (n=44) (all drawn from NEPTUNE (56)) and AAV (n=15) (drawn from ERCB) (28), accessed in GEO by accession GSE108113. Differential expression analysis was restricted to probe sets (n=12,074) common to both Affymetrix microarray platforms (GeneChip Human Genome U133A 2.0 and U133 Plus 2.0, ThermoFisher) by comparing transcriptional profiles from individuals with CKD versus living donors using the SAM (significance analysis of microarrays) method (28) and genes were defined as differentially expressed if they met q-value < 0.05.

Aggregate gene expression scores (Fig. 5A) were calculated based on the genes from Table S1 for “EGE” (C1) and “MPC” (C7). Genes shared by these two lists were used to generate a “Shared” gene list, with the residual genes in each of the EPC and MPC lists were used to define those clusters. Gene lists were further distilled to the subset of genes queried on the microarray. An aggregate gene expression score was then generated for each of these gene lists by averaging expression of all genes, after each gene’s log2 expression was Z-score transformed.

GFR in ERCB and NEPTUNE was estimated by the four-variable Modification of Diet in Renal Disease (MDRD) study equation (57) and log2 transformed prior to correlation with expression data as described (28). eGFRs were available in all 191 individuals in the discovery cohort and 101 individuals in the validation cohort. Correlation analyses between gene expression and log2 eGFR were performed using Pearson’s correlation coefficient.

Correlation of gene expression with proteinuria was performed in the ERCB lupus nephritis cohort in which proteinuria was quantified (58) (GEO accession GSE32591), accessed in Nephroseq (www.nephroseq.org, Berthier Lupus Glomerulonephritis dataset) and analyzed using the inherent analysis mechanism which includes pre-computation of differential expression profiles using Student’s t-test for two class differential expression analyses. In the validation cohort, 88 individuals had a urine protein to creatinine ratio (UPCR) available, and did not include individuals with AAV.

### Statistics

Methods of statistical analysis are included in the relevant Methods sections and figure legends.

### Study and ethics approval

Study approval was granted by the University of Michigan Human Pluripotent Stem Cell Research Oversight compliance program (Application #1096), and the ethics committees for ERCB and NEPTUNE studies. The University of Michigan Institutional Review Board approved gene expression studies (HUM0002468), and human kidney tissue procurement (HUM00045864, HUM00083116). Written informed consent was received from participants prior to inclusion.

### Data Access

Normalized organoid scRNA-seq gene expression data files reported herein were submitted to NCBI Gene Expression Omnibus, GEO Series accession number GSE115986.

## AUTHOR CONTRIBUTIONS

All authors critically reviewed the manuscript. JLH planned and performed organoid experiments, supervised NLW, designed and managed the project, critically reviewed data, and wrote the manuscript. NLW and EAO planned and performed organoid and Drop-seq experiments, respectively. NLW performed immunostaining of human kidney tissue. RM performed single cell transcriptomic analysis for organoids and developing human kidney. JZ performed trajectory analysis of single cell transcriptomes. SE analyzed human tissue transcriptome-disease correlations and identified high priority genes. VN analyzed rat transcriptomic data. CC and JRS provided developing human kidney data and provided expertise related to stem cell culture protocols. OGT supervised JZ. JBH and RCW provided and analyzed human kidney tissue samples. RCW also provided rat tissue transcriptomic data and critically reviewed data. BSF provided organoid and stem cell expertise, and critically reviewed data. MK critically reviewed data, and supervised the entire project.

## ACKNOWLEDGEMENTS

The authors extend their gratitude to investigators and participants involved in the ERCB and the NEPTUNE consortium (detailed in Supplemental Acknowledgments). This research was supported by NIH grants: K08 DK089119 to JLH, U54 DK083912 to MK, and P30 DK081943 Applied Systems Biology Core at the University of Michigan George M O’Brien Kidney Translational Core Center and P30 CA046592 Rogel Cancer Center Tissue and Molecular Pathology Shared Resource. Additional funding and/or programmatic support was provided by the NephCure Kidney International and the Halpin Foundation through support for NEPTUNE.

## Notes

Conflict of interest statement: The authors have declared that no conflict of interest exists

## REFERENCES

1. Liu Z-H, and He JC. Podocytopathy. Basel: Karger; 2014.

2. Hodgin JB, Nair V, Zhang H, Randolph A, Harris RC, Nelson RG, et al. Identification of cross-species shared transcriptional networks of diabetic nephropathy in human and mouse glomeruli. Diabetes. 2013;62(1):299–308.

3. Shankland SJ, Pippin JW, Reiser J, and Mundel P. Podocytes in culture: past, present, and future. Kidney international. 2007;72(1):26–36.

4. Cohen CD, Frach K, Schlondorff D, and Kretzler M. Quantitative gene expression analysis in renal biopsies: a novel protocol for a high-throughput multicenter application. Kidney international. 2002;61(1):133–40.

5. Cohen CD, Grone HJ, Grone EF, Nelson PJ, Schlondorff D, and Kretzler M. Laser microdissection and gene expression analysis on formaldehyde-fixed archival tissue. Kidney international. 2002;61(1):125–32.

6. Bennett MR, Czech KA, Arend LJ, Witte DP, Devarajan P, and Potter SS. Laser capture microdissection-microarray analysis of focal segmental glomerulosclerosis glomeruli. Nephron Exp Nephrol. 2007;107(1):e30–40.

7. Takemoto M, He L, Norlin J, Patrakka J, Xiao Z, Petrova T, et al. Large-scale identification of genes implicated in kidney glomerulus development and function. The EMBO journal. 2006;25(5):1160–74.

8. Ju W, Greene CS, Eichinger F, Nair V, Hodgin JB, Bitzer M, et al. Defining cell-type specificity at the transcriptional level in human disease. Genome Res. 2013;23(11):1862–73.

9. Freedman BS, Brooks CR, Lam AQ, Fu H, Morizane R, Agrawal V, et al. Modelling kidney disease with CRISPR-mutant kidney organoids derived from human pluripotent epiblast spheroids. Nat Commun. 2015;6:8715.

10. Takasato M, Er PX, Chiu HS, Maier B, Baillie GJ, Ferguson C, et al. Kidney organoids from human iPS cells contain multiple lineages and model human nephrogenesis. Nature. 2015;526(7574):564–8.

11. Morizane R, Lam AQ, Freedman BS, Kishi S, Valerius MT, and Bonventre JV. Nephron organoids derived from human pluripotent stem cells model kidney development and injury. Nat Biotechnol. 2015;33(11):1193–200.

12. Taguchi A, Kaku Y, Ohmori T, Sharmin S, Ogawa M, Sasaki H, et al. Redefining the in vivo origin of metanephric nephron progenitors enables generation of complex kidney structures from pluripotent stem cells. Cell stem cell. 2014;14(1):53–67.

13. Macosko EZ, Basu A, Satija R, Nemesh J, Shekhar K, Goldman M, et al. Highly Parallel Genome-wide Expression Profiling of Individual Cells Using Nanoliter Droplets. Cell. 2015;161(5):1202–14.

14. Adam M, Potter AS, and Potter SS. Psychrophilic proteases dramatically reduce single-cell RNA-seq artifacts: a molecular atlas of kidney development. Development. 2017;144(19):3625–32.

15. Czerniecki SM, Cruz NM, Harder JL, Menon R, Annis J, Otto EA, et al. High-Throughput Screening Enhances Kidney Organoid Differentiation from Human Pluripotent Stem Cells and Enables Automated Multidimensional Phenotyping. Cell stem cell. 2018;22(6):929–40 e4.

16. Cruz NM, Song X, Czerniecki SM, Gulieva RE, Churchill AJ, Kim YK, et al. Organoid cystogenesis reveals a critical role of microenvironment in human polycystic kidney disease. Nature materials. 2017;16(11):1112–9.

17. Kim YK, Refaeli I, Brooks CR, Jing P, Gulieva RE, Hughes MR, et al. Gene-Edited Human Kidney Organoids Reveal Mechanisms of Disease in Podocyte Development. Stem cells. 2017;35(12):2366–78.

18. Uhlen M, Fagerberg L, Hallstrom BM, Lindskog C, Oksvold P, Mardinoglu A, et al. Proteomics. Tissue-based map of the human proteome. Science. 2015;347(6220):1260419.

19. Little MH, and ScienceDirect (Online service). Amsterdam; Boston: Elsevier/AP, Academic Press is an imprint of Elsevier; 2016.

20. Whitfield ML, George LK, Grant GD, and Perou CM. Common markers of proliferation. Nature reviews Cancer. 2006;6(2):99–106.

21. Menon R, Otto EA, Kokoruda A, Zhou J, Zhang Z, Yoon E, et al. Single-cell analysis of progenitor cell dynamics and lineage specification of the human fetal kidney. bioRxiv. 2018.

22. Lambert SA, Jolma A, Campitelli LF, Das PK, Yin Y, Albu M, et al. The Human Transcription Factors. Cell. 2018;172(4):650–65.

23. Sonawane AR, Platig J, Fagny M, Chen CY, Paulson JN, Lopes-Ramos CM, et al. Understanding Tissue-Specific Gene Regulation. Cell reports. 2017;21(4):1077–88.

24. Carro MS, Lim WK, Alvarez MJ, Bollo RJ, Zhao X, Snyder EY, et al. The transcriptional network for mesenchymal transformation of brain tumours. Nature. 2010;463(7279):318–25.

25. Walsh LA, Alvarez MJ, Sabio EY, Reyngold M, Makarov V, Mukherjee S, et al. An Integrated Systems Biology Approach Identifies TRIM25 as a Key Determinant of Breast Cancer Metastasis. Cell reports. 2017;20(7):1623–40.

26. Motojima M, Kume T, and Matsusaka T. Foxc1 and Foxc2 are necessary to maintain glomerular podocytes. Experimental cell research. 2017;352(2):265–72.

27. Maezawa Y, Onay T, Scott RP, Keir LS, Dimke H, Li C, et al. Loss of the podocyte-expressed transcription factor Tcf21/Pod1 results in podocyte differentiation defects and FSGS. Journal of the American Society of Nephrology: JASN. 2014;25(11):2459–70.

28. Grayson PC, Eddy S, Taroni JN, Lightfoot YL, Mariani L, Parikh H, et al. Metabolic pathways and immunometabolism in rare kidney diseases. Annals of the rheumatic diseases. 2018;77(8):1226–33.

29. Tao J, Mariani L, Eddy S, Maecker H, Kambham N, Mehta K, et al. JAK-STAT signaling is activated in the kidney and peripheral blood cells of patients with focal segmental glomerulosclerosis. Kidney international. 2018.

30. Cho EA, Patterson LT, Brookhiser WT, Mah S, Kintner C, and Dressler GR. Differential expression and function of cadherin-6 during renal epithelium development. Development. 1998;125(5):803–12.

31. Menon R, Otto EA, Kokoruda A, Zhou J, Zhang Z, Yoon E, et al. Single-cell analysis of progenitor cell dynamics and lineage specification of the human fetal kidney. Development. In press.

32. Nishizono R, Kikuchi M, Wang SQ, Chowdhury M, Nair V, Hartman J, et al. FSGS as an Adaptive Response to Growth-Induced Podocyte Stress. Journal of the American Society of Nephrology: JASN. 2017;28(10):2931–45.

33. Chen IH, Wang HH, Hsieh YS, Huang WC, Yeh HI, and Chuang YJ. PRSS23 is essential for the Snail-dependent endothelial-to-mesenchymal transition during valvulogenesis in zebrafish. Cardiovascular research. 2013;97(3):443–53.

34. Gharib SA, Pippin JW, Ohse T, Pickering SG, Krofft RD, and Shankland SJ. Transcriptional landscape of glomerular parietal epithelial cells. PLoS One. 2014;9(8):e105289.

35. Smeets B, Angelotti ML, Rizzo P, Dijkman H, Lazzeri E, Mooren F, et al. Renal progenitor cells contribute to hyperplastic lesions of podocytopathies and crescentic glomerulonephritis. Journal of the American Society of Nephrology: JASN. 2009;20(12):2593–603.

36. Kuppe C, Grone HJ, Ostendorf T, van Kuppevelt TH, Boor P, Floege J, et al. Common histological patterns in glomerular epithelial cells in secondary focal segmental glomerulosclerosis. Kidney international. 2015;88(5):990–8.

37. Smeets B, Stucker F, Wetzels J, Brocheriou I, Ronco P, Grone HJ, et al. Detection of activated parietal epithelial cells on the glomerular tuft distinguishes early focal segmental glomerulosclerosis from minimal change disease. Am J Pathol. 2014;184(12):3239–48.

38. Appel D, Kershaw DB, Smeets B, Yuan G, Fuss A, Frye B, et al. Recruitment of podocytes from glomerular parietal epithelial cells. Journal of the American Society of Nephrology: JASN. 2009;20(2):333–43.

39. Grgic I, Hofmeister AF, Genovese G, Bernhardy AJ, Sun H, Maarouf OH, et al. Discovery of new glomerular disease-relevant genes by translational profiling of podocytes in vivo. Kidney international. 2014;86(6):1116–29.

40. Eng DG, Kaverina NV, Schneider RRS, Freedman BS, Gross KW, Miner JH, et al. Detection of renin lineage cell transdifferentiation to podocytes in the kidney glomerulus with dual lineage tracing. Kidney international. 2018;93(5):1240–6.

41. Guhr SS, Sachs M, Wegner A, Becker JU, Meyer TN, Kietzmann L, et al. The expression of podocyte-specific proteins in parietal epithelial cells is regulated by protein degradation. Kidney international. 2013;84(3):532–44.

42. Park J, Shrestha R, Qiu C, Kondo A, Huang S, Werth M, et al. Single-cell transcriptomics of the mouse kidney reveals potential cellular targets of kidney disease. Science. 2018;360(6390):758–63.

43. Wu H, Uchimura K, Donnelly E, Kirita Y, Morris SA, and Humphreys BD. Comparative analysis of kidney organoid and adult human kidney single cell and single nucleus transcriptomes. bioRxiv. 2017.

44. Combes AN, Phipson B, Zappia L, Lawlor K, Er PX, Oshlack A, et al. High throughput single cell RNA-seq of developing mouse kidney and human kidney organoids reveals a roadmap for recreating the kidney. bioRxiv. 2017.

45. Barthel LK, and Raymond PA. Improved method for obtaining 3-microns cryosections for immunocytochemistry. The journal of histochemistry and cytochemistry: official journal of the Histochemistry Society. 1990;38(9):1383–8.

46. Venkatareddy M, Wang S, Yang Y, Patel S, Wickman L, Nishizono R, et al. Estimating podocyte number and density using a single histologic section. Journal of the American Society of Nephrology: JASN. 2014;25(5):1118–29.

47. Yang Y, Hodgin JB, Afshinnia F, Wang SQ, Wickman L, Chowdhury M, et al. The two kidney to one kidney transition and transplant glomerulopathy: a podocyte perspective. Journal of the American Society of Nephrology: JASN. 2015;26(6):1450–65.

48. Kim YH, Goyal M, Kurnit D, Wharram B, Wiggins J, Holzman L, et al. Podocyte depletion and glomerulosclerosis have a direct relationship in the PAN-treated rat. Kidney international. 2001;60(3):957–68.

49. Butler A, Hoffman P, Smibert P, Papalexi E, and Satija R. Integrating single-cell transcriptomic data across different conditions, technologies, and species. Nat Biotechnol. 2018;36(5):411–20.

50. Shekhar K, Lapan SW, Whitney IE, Tran NM, Macosko EZ, Kowalczyk M, et al. Comprehensive Classification of Retinal Bipolar Neurons by Single-Cell Transcriptomics. Cell. 2016;166(5):1308–23 e30.

51. Schmid H, Cohen CD, Henger A, Schlondorff D, and Kretzler M. Gene expression analysis in renal biopsies. Nephrology, dialysis, transplantation: official publication of the European Dialysis and Transplant Association - European Renal Association. 2004;19(6):1347–51.

52. Gadegbeku CA, Gipson DS, Holzman LB, Ojo AO, Song PX, Barisoni L, et al. Design of the Nephrotic Syndrome Study Network (NEPTUNE) to evaluate primary glomerular nephropathy by a multidisciplinary approach. Kidney international. 2013;83(4):749–56.

53. Levey AS, Coresh J, Greene T, Stevens LA, Zhang YL, Hendriksen S, et al. Using standardized serum creatinine values in the modification of diet in renal disease study equation for estimating glomerular filtration rate. Annals of internal medicine. 2006;145(4):247–54.

54. Berthier CC, Bethunaickan R, Gonzalez-Rivera T, Nair V, Ramanujam M, Zhang W, et al. Cross-species transcriptional network analysis defines shared inflammatory responses in murine and human lupus nephritis. Journal of immunology. 2012;189(2):988–1001.

